# Synchronous diversification of Sulawesi’s iconic artiodactyls driven by recent geological events

**DOI:** 10.1101/241448

**Authors:** Laurent A. F. Frantz, Anna Rudzinski, Abang Mansyursyah Surya Nugraha, Allowen Evin, James Burton, Ardern Hulme-Beaman, Anna Linderholm, Ross Barnett, Rodrigo Vega, Evan K. Irving-Pease, James Haile, Richard Allen, Kristin Leus, Jill Shephard, Mia Hillyer, Sarah Gillemot, Jeroen van den Hurk, Sharron Ogle, Cristina Atofanei, Mark G. Thomas, Friederike Johansson, Abdul Haris Mustari, John Williams, Kusdiantoro Mohamad, Chandramaya Siska Damayanti, Ita Djuwita Wiryadi, Dagmar Obbles, Stephano Mona, Hally Day, Muhammad Yasin, Stefan Meker, Jimmy A. McGuire, Ben J. Evans, Thomas von Rintelen, Simon Y. W. Ho, Jeremy B. Searle, Andrew C. Kitchener, Alastair A. Macdonald, Darren J. Shaw, Robert Hall, Peter Galbusera, Greger Larson

**Author notes:** deceased (Ita Djuwita Wiryadi). contributed equally (Laurent A. F. Frantz, Anna Rudzinski, Abang Mansyursyah Surya Nugraha, Allowen Evin, James Burton). co-supervised the study (Alastair A. Macdonald, Darren J. Shaw, Robert Hall, Peter Galbusera, Greger Larson). Present address: Pertamina University, Jl. Teuku Nyak Arief, Kawasan Simprug, Kebayoran Lama, Jakarta Selatan 12220, Indonesia.

## Abstract

The high degree of endemism on Sulawesi has previously been suggested to have vicariant origins, dating back 40 Myr ago. Recent studies, however, suggest that much of Sulawesi’s fauna assembled over the last 15 Myr. Here, we test the hypothesis that recent uplift of previously submerged portions of land on Sulawesi promoted diversification, and that much of the its faunal assemblage is much younger than the island itself. To do so, we combined palaeogeographical reconstructions with genetic and morphometric data sets derived from Sulawesi’s three largest mammals: the Babirusa, Anoa, and Sulawesi warty pig. Our results indicate that although these species most likely colonized the area that is now Sulawesi at different times (14 Myr ago to 2-3 Myr ago), they experienced an almost synchronous expansion from the central part of the island. Geological reconstructions indicate that this area was above sea level for most of the last 4 Myr, unlike most parts of the island. We conclude that recent emergence of land on Sulawesi (~1–2 Myr) may have allowed species to expand synchronously. Altogether, our results indicates that the establishment of the highly endemic faunal assemblage on Sulawesi was driven by geological events over the last few million years.

## Introduction

Alfred Russel Wallace was the first to document the ‘anomalous’ biogeographic region in Island Southeast Asia now known as Wallacea [1,2]. This biodiversity hotspot [3] is bounded by Wallace’s Line in the west and Lydekker’s Line in the east [4]. It consists of numerous islands in the Indonesian archipelago, all of which boast a high degree of endemism. For example, on Sulawesi, the largest island in the region, at least 61 non-volant mammalian species are endemic [5] and this figure is likely to be an underestimate.

The geological origins of Wallacea are as complex as its biogeography. Until recently, Sulawesi had been regarded as the product of multiple collisions of continental fragments from the Late Cretaceous [6–9]. This assumption has been challenged and a recent reinterpretation suggests instead that the island began to form as the result of continental collisions during the Cretaceous, which were then followed by Eocene rifting of the Makassar Strait. This process led to the isolation of small land areas in western Sulawesi from Sundaland. In the early Miocene, a collision with the Australian crust of the Sula Spur led to l uplift and emergence of land [10–12]. Later extension, uplift and subsidence from the middle Miocene to the present day led to the modern uneven distribution of micro-continental fragments, and the emergence of islands (separated by deep seas) between Borneo and Australia [13,14].

Previous geological interpretation involving the assembly of multiple terranes by collision was used to suggest that Sulawesi’s peculiar species richness resulted from vicariance and amalgamation over long geological time periods [10,15,16]. However, recent molecular-clock analyses suggest that a dispersal, starting in the middle Miocene (~15 Myr ago) from both Sunda and Sahul is a more plausible explanation [17,18] [17,19][17,18]. These conclusions suggest a limited potential for animal dispersal to Sulawesi prior to ~15 Myr ago. Rapid tectonic changes, coupled with the dramatic sea-level fluctuations over the past 5 Myr [20], might also have affected land availability and influenced patterns of species dispersal to Sulawesi, intra-island species expansion and speciation.

The hypothesis of a recent increase in land area [19] can be tested by comparing the population histories of multiple species on the island. Analyses of genetic and morphometric variability can be used to infer the timing and trajectories of dispersal, and the geographical and temporal origins of expansion. For example, if land area had increased, from a single smaller island, extant species now living on Sulawesi, would all have expanded from the same area. In addition, under this assumption, within the same geographical region their respective diversifications would be expected to have been roughly simultaneous.

Here, we focus on three large mammals endemic to Sulawesi: the Babirusa (*Babyrousa* spp.), the Sulawesi warty pig (SWP, *Sus celebensis*) and the Anoa, a dwarf buffalo, (*Bubalus* spp.). The Babirusa (*Babyrousa spp*.) is a suid characterized by wrinkled skin and four extraordinary curved canine tusks displayed by males [21–23]. It represents a “ghost lineage” since there are no closely related extant species outside Sulawesi (*e.g*. African suids are more closely related to the Babirusa than Asian suids) and the Babirusa is unknown in the fossil record outside Sulawesi [24]. Three extant species of Babirusa (distributed primarily in the interior of Sulawesi and on surrounding islands [21–23] have been described: *Babyrousa babyrousa* (Buru and Sulu islands), *Babyrousa celebensis* (mainland Sulawesi) and *Babyrousa togeanensis* (Togian Island) [25].

The Anoa is an endemic “miniature buffalo” related to indigenous bovids in the Philippines and East Asia [26]. It stands approximately one metre tall, weighs 150–200 kg, and mostly inhabits pristine rainforest [27]. Although the subgenus *Anoa* has been divided into two species, the lowland Anoa (*Bubalus depressicornis*) and the highland Anoa (*Bubalus quarlesi*) [28], this classification is still contentious [29]. In contrast with Anoa and Babirusa, the Sulawesi warty pig (SWP; *Sus celebensis*) occupies a wide range of habitats, ranging from swamps to rainforests. This species is closely related to the Eurasian wild pig(Sus *scrofa*), from which it diverged during the early Pleistocene (~2 Myr ago)[24,30]. The SWP has been found on numerous islands throughout Island Southeast Asia (ISEA), probably as the result of human-mediated dispersal [31]. As its name implies, male SWPs develop facial warts. These three cultural icons (represented by some of the oldest prehistoric cave paintings [32,33]) have undergone recent and significant population reduction and range contraction due to overhunting and conversion of natural habitat for agricultural use.

Here, we establish when Sulawesi gained its modern shape and size, including connectivity between its constituent peninsulae, and assessed the impact of island formation on the evolution of Sulawesi’s biodiversity. To do so, we used new reconstructions of the island’s palaeogeography that allowed us to interpret the distribution of land and sea over the last 8 Myr at 1 Myr intervals. To determine the timings of diversification of the three largest endemic mammals on the island, we generated and analysed genetic and morphometric data from a total of 1,289 samples of the SWP, Anoa, and Babirusa obtained from museums, zoos and wild populations (456, 520 and 313 samples respectively; Table S1). Although these taxa have been divided into multiple species (see taxonomic notes in the Supplementary Material), for the purpose of this study we treated SWP, Anoa and Babirusa as single taxonomic units.

## Results and Discussion

### Contemporaneous divergence

We generated mitochondrial DNA (mtDNA) sequences and/or microsatellite data from 230 SWPs, 155 Anoas and 213 Babirusas sampled across Sulawesi and the neighbouring islands (Supplementary Material Figure S1; Table S1). Using a molecular-clock analysis, we inferred the time to the most recent common ancestor (TMRCA) of each species. The estimates from this method represent coalescence times, which provide a reflection of the crown age of each taxon. The closer relationship between Babirusa and SWP (~13 Myr ago) [34], compared with the divergence of either species from the Anoa (~58 Myr ago)[35] allowed us to align sequences from Babirusa and SWP alongside one another and jointly infer their relative TMRCAs. Separate analyses were performed for the Anoa. The inferred TMRCA of SWP was 2.19 Myr (95% credibility interval [CI] 1.19–3.41 Myr; Supplementary Material Figure S2) and Babirusa was 2.49 Myr (95% CI 1.33–3.61 Myr) (Figure 1a; Supplementary Material Figure S2). The inferred TMRCA of Anoa was younger (1.06 Myr; Figure 1a; Supplementary Material Figure S3), though its 95% CI (0.81–1.96 Myr) overlapped substantially with the TMRCAs of the other two species.

**Figure 1:**
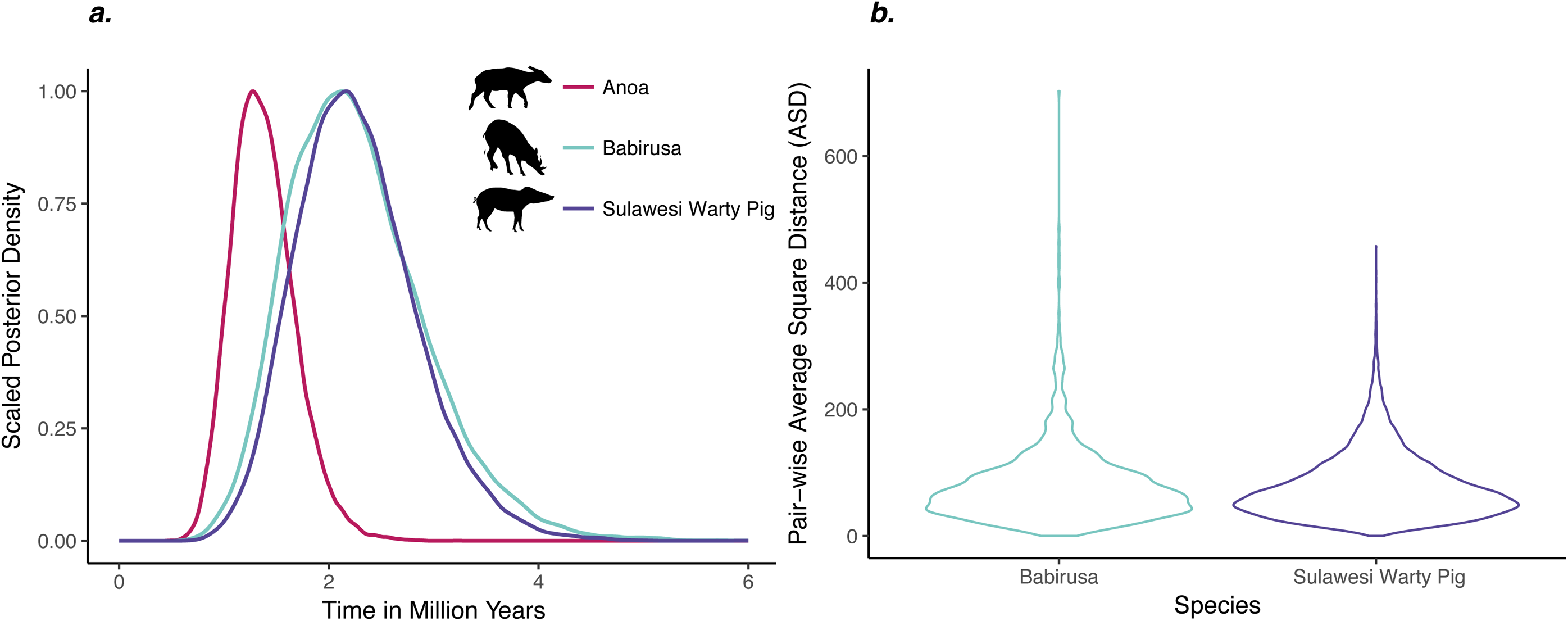
Time to the most recent common ancestor (TMRCA) for three mammal species on Sulawesi. **a**. Posterior densities of the TMRCA estimates for Anoa, Babirusa, and Sulawesi warty pig inferred using a Bayesian molecular clock based on mitochondrial DNA sequences. **b**. Distribution of the average squared distances computed from microsatellite data from Babirusa and Sulawesi warty pig.

The relatively recent divergence between Babirusa and SWP also allowed us compare their TMRCAs using identical microsatellite loci. To do so, we computed the average square distance (ASD)[36,37] between every pair of individuals within each species at the same 13 microsatellite loci. Although such an analysis might be affected by population structure (see below), we found that the distributions of ASD values were not significantly different between these two species (Wilcoxon signed-rank test, *p*=0.492; Figure 1b). This is consistent with the mitochondrial evidence for the nearly identical TMRCAs in the two species.

Recent molecular analyses have indicated that Babirusa may have colonized Wallacea as early as 13 Myr ago, whereas SWP and Anoa appear to have only colonized Sulawesi within the last 2–4 Myr [17,30,32,34]. An early dispersal of Babirusa to Sulawesi (late Palaeogene) has also been suggested on the basis of palaeontological evidence [19]. In addition, our data corroborate previous studies in indicating that both SWP and Babirusa are monophyletic with respect to their most closely related taxa on neighbouring islands (*e.g*. Borneo), which is consistent with only one colonization of Sulawesi (Supplementary Material; Figure S4-6)[30].

We then examined whether patterns of morphological diversity in these taxa are consistent with the molecular date estimates. To do so, we obtained measurements of 356 second and third lower molar (M2 and M3) from 95 Babirusas and 132 SWPs. SWP and Babirusa do not overlap morphologically (Figure 2a) and we were thus able to assign each specimen to its correct species with success rates of 94.3% (CI: 92.7%–95.5%, distribution of leave-one-out cross validation of a discriminant analysis based on a balanced sample design) [38] and 94.7% (CI: 93.8%–96.7%) based on their M2 and M3, respectively. Our results also indicate that Babirusa did not accumulate more tooth shape variation within Sulawesi (Fligner-Killeen test X^2^=1.04, *p*=0.3 for M2, X^2^=3.45, *p*=0.06 for M3). The data instead suggests that SWP has greater variance in the size of its M3 (X^2^=4.52, *p*=0.03, but not in the size of the M2, X^2^=3.44, *p*=0.06), and that the population from West Central Sulawesi has an overall smaller tooth size than the two populations from North West and North East Sulawesi (Figure 2b, Table S2). While these results may result from different selective constraints, they indicate that Babirusa did not accumulate greater morphological variation in tooth shape than did the SWP, despite arriving on Sulawesi up to 10 Myr earlier.

**Figure 2:**
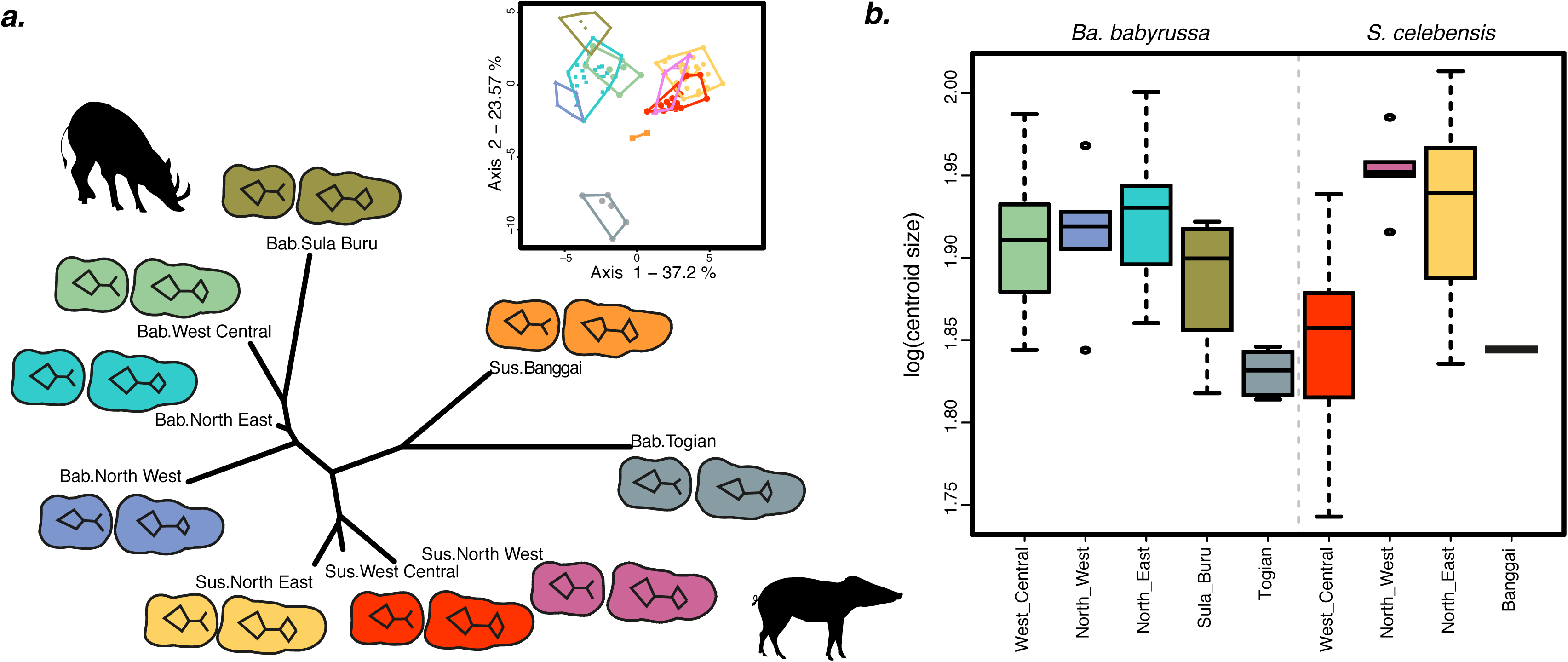
Population morphological variation inferred from geometric morphometric data. **a**. Neighbour-joining network based on Mahalanobis distances measured from second and third lower molar shapes and visualisation of population mean shape. **b**. Variation of third molar size per population (log centroid size).

Altogether our analyses suggest that although the three species are believed to have colonized the island at different times, their similar degrees of morphological diversity and their nearly synchronous TMRCAs raise the possibility that they (and possibly other species) responded to a common mechanism that triggered their contemporaneous diversification.

### Past land availability correlates with the expansion origins

Increasing land area may have promoted a simultaneous diversification and range expansion in Babirusas, SWPs, and Anoas. To test this hypothesis, we used a new reconstruction that depicts land area in the Sulawesi region through time using information from the geological record. The reconstructions in 1 Myr increments (Figure 3a; Figure S7; [39]) support a scenario in which most of Sulawesi was submerged until the late Pliocene to early Pleistocene (2–3 Myr ago). Large-scale uplifts over the last 2–3 Myr would have rapidly and significantly increased land area, making it possible for non-volant species to expand their ranges.

**Figure 3:**
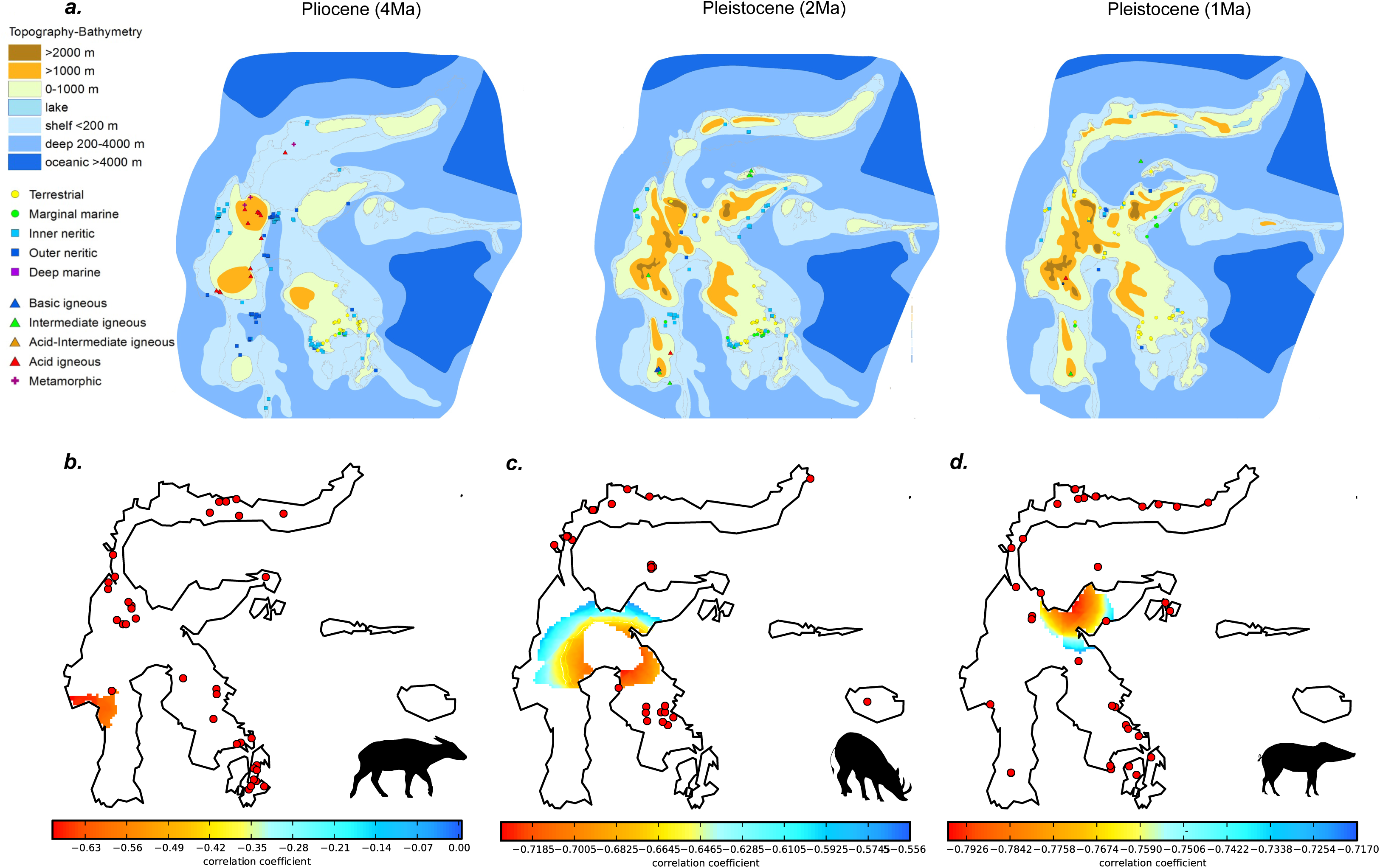
Geological maps of Sulawesi and the geographical origin of expansion. **a**. Reconstruction of Sulawesi over the last 5 Myr (adapter from [39]) and potential origin of expansion of **b**. Anoa, **c**. Babirusa, and **d**. Sulawesi warty pig. Low correlation values (between distance and extrapolated genetic diversity; see *Supplementary Material*) represent most likely origin of expansion.

To further assess whether these Plio-Pleistocene uplifts were responsible for a synchronous expansion, we inferred the most likely geographical origin of expansion using microsatellite data under a model of spatial loss-of-diversity with distance from expansion origin (Supplementary Material). These estimates were obtained independently of, and uninformed by, either the geological reconstructions or modern phylogeographical boundaries inferred from other species. We deduced that the most likely origin for both SWP and Babirusa was in the East Central region of Sulawesi (Figure 3c and 3d), and the most likely origin of Anoa was in the West Central region (Figure 3b).

The origins of the population expansions of both SWP and Babirusa occurred in an area of Sulawesi that only emerged during the late Pliocene to early Pleistocene (Figure 3a; Supplementary Material Figure S7). On the other hand, the Anoa most likely origin of diversification lies in a region that was submerged until the Pleistocene, consistent with paleontological evidence [32] and with the slightly more recent TMRCA inferred for this species (Figure 1a). Thus, for all three species, the inferred geographical origins of their range expansions match the land availability derived from our geological reconstruction of Sulawesi.

### Geological history of past land isolation correlates with zones of endemism

Previous studies have identified endemic zones that are common to macaques, toads [18,40], tarsiers [41–44] and lizards [45]. We tested whether the same area of endemism are linked to the population structure in our three species by generating a phylogenetic tree for each species using mtDNA and defined 5–6 haplogroups per species based on well-supported clades (Figure 4a-c; *Supplementary Material* Figure S4-6). We found that haplogroup proportions were significantly different between previously defined areas of endemism in all three species (Pearson’s chi-squared test; *p*<0.001), suggesting population substructure.

**Figure 4:**
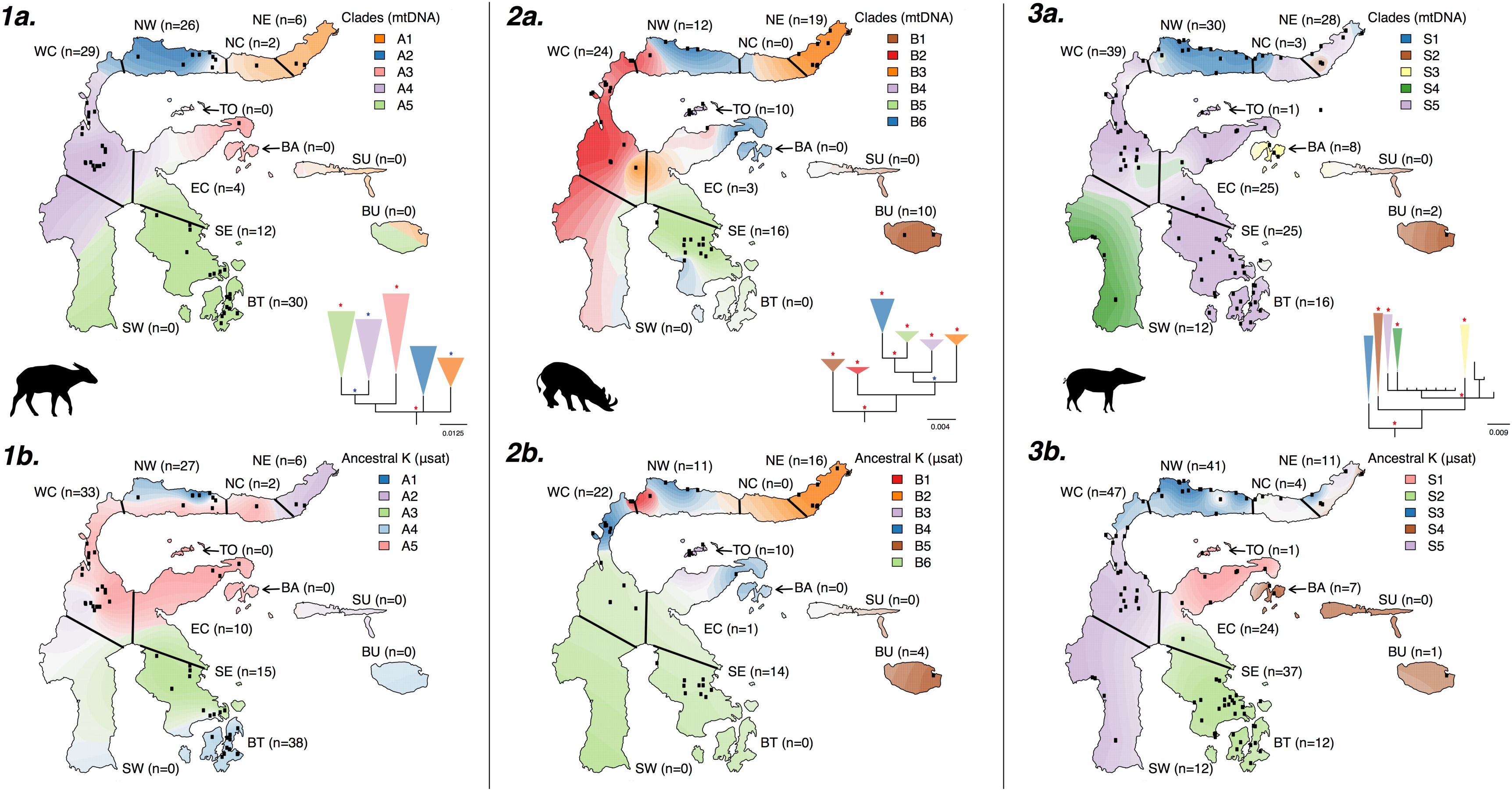
Population structure and geographic patterning of three mammal species on Sulawesi inferred from mitochondrial and microsatellite DNA. **a**. to **c**., A tessellated projection of sample haplogroups in each region of endemism, and phylogeny of **a**. Anoa **b**. Babirusa, and **c**. Sulawesi warty pig. Each region is labelled with the number of samples used for the projection. The projection extends over regions with no samples (*e.g*. the Southwest peninsula for Babirusa and Anoa) and the population membership affinities for these regions are therefore less reliable. Red and blue stars on the phylogenetic trees correspond to posterior probabilities greater than 0.9 and 0.7, respectively. **d. to f**., Tessellated projection of the STRUCTURE analysis, using the microsatellite data, for **d**. Anoa, **e**. Babirusa, and **f**. Sulawesi warty pig. The best *K* value for each species was used (*K*=5 for Anoa; *K*=6 for Babirusa; *K*=5 for Sulawesi warty pig; *Supplementary Material* Figure S8). NE=North East; NC=North Central; NW=North West; TO=Togian; BA=Banggai Archipelago; EC=East Central; WC=West Central; SU=Sula; BU=Buru; SE=South East; SW= South West; BT=Buton.

We also used STRUCTURE [46] to infer population structure from microsatellite data. The optimum numbers of populations (*K*) were 5, 6 and 5 for Anoa, Babirusa and SWP, respectively (Supplementary Material Figure S8; Figure 4d-f). Plotting the proportion of membership of each sample onto a map revealed a strong correspondence with the previously described zones of endemism (Figure 4d-f). Using an analysis of molecular variance (AMOVA), we found that these areas of endemism explained approximately 17%, 27%, and 5% of the variance in allele frequencies in Anoa, Babirusa and SWP, respectively (Table S5). Populations of Babirusa and SWP in these zones of endemism were also strongly morphologically differentiated (Figure 2).

Altogether, these data and analyses indicate that, despite some differences, the zones of endemism identified in tarsiers, macaques, toads and lizards [18,40–45,47] are largely consistent with the population structure and morphological differentiation in the three species studied here. This is particularly striking for the north arm of Sulawesi (NW, NC, and NE in Figure 4), where we identify two highly differentiated populations (reflected in both mtDNA and nuclear data sets) in all three taxa. This pattern could result from either adaptation to local environments or from isolation due to the particular geological history associated with the northern arm. Geological reconstructions (Figure 3a) indicate that although land was present in this region during the past 4 Myr, it was often isolated from the rest of Sulawesi until the mid-Pleistocene. Thus, the combined geological and biological evidence presented here indicate that the high degree of divergence observed in the northern-arm populations in a multitude of species (*e.g*. three ungulates, macaques, and tarsiers) might have been shaped by isolation from the rest of the island until until the last 1My (Figure 3a).

### Recent and contemporary land isolation also affected morphological evolution including dwarfism

Similar isolation is likely to have influenced the populations inhabiting the smaller islands adjacent to Sulawesi, including the Banggai archipelago, Buru, the Togian and Sula Islands. Interestingly, our geometric morphometric analyses demonstrated that these island populations of SWP and Babirusa are the most morphologically divergent (Figure 2a). For example, the insular populations from the Togian Islands (Babirusa) and the Banggai archipelago (SWP) were found to have much smaller tooth sizes than their counterparts on the mainland (Figure 2b).

The significant morphometric divergences between populations on various islands are consistent with the genetic differentiation between Babirusa/SWP on Togian, Sula, and Buru (Figure 4; *Supplementary Material* Figure S9; *Supplementary Material* Figure S10) and between island populations of SWP on Banggai archipelago, Buton, and Buru (Figure 4; *Supplementary Material* Figure S9; *Supplementary Material* Figure S10).

Together, these results show that while suture zones between tectonic fragments are consistent with genetic and morphometric differentiation within Sulawesi, isolation on remote islands is likely to have had a much greater effect on morphological distinctiveness. Rapid evolution, on islands, has been described in many species (e.g [48]). Dwarfism on small islands has previously been suggested to be the result of environmental constraints, including in pigs [49] where island populations are known to have smaller tooth sizes than their mainland counterparts [50,51].

### Demographic history

Isolation of subpopulations across Sulawesi might also be linked to recent anthropogenic disturbances, especially for Anoa and Babirusa, that occupy pristine forest or swamps [21,27]. In order to assess the impact of recent anthropogenic changes on the three species, we inferred their demographic history using approximate Bayesian computation (ABC). We fitted various demographic models to the genetic data (combining both mtDNA and microsatellite data; *Supplementary Material;* Figure S11). The best-supported demographic model involved a long-term expansion followed by a recent bottleneck in all three species (Table S3), corroborating the results of recent analyses of the SWP genome [30].

While our ABC analysis had insufficient power to retrieve the time of expansion (Table S4), it provided relatively narrow estimates of the current effective population sizes (Figure 5; Table S4). We inferred a larger effective population size in SWP (83,021; 95% CI 46,287–161,457) than in Babirusa (30,895; 95% CI 17,522–54,954) or Anoa (27,504; 95% CI 13,680–54,056). *Sus celebensis* occupies a wide range of habitats, including agricultural areas [52]. Thus, this species is likely to be less affected by continuing deforestation than Babirusa or Anoa, which are typically restricted to less disturbed forest and swamps [21,26]. Phylogenetic analyses of microsatellite data indicate more geographical structuring in Babirusa and Anoa than in SWP (*Supplementary Material* Figure S12; Table S5). Overall, our results are consistent with habitat loss and fragmentation coupled with species-specific differences in habitat tolerance have affected population size and structure.

**Figure 5:** Posterior distribution of the current population size (Ne) of each species as inferred via approximate Bayesian computation.

## Conclusions

Our results indicate that, while the different geological component of Sulawesi were assembled at about 23 Myr ago, the island only acquired its distinctive modern form in the last few million years. By 3 Myr ago there was a large single island at its modern centre, but the complete connection between the arms was established more recently. The increasing land area associated with Plio-Pleistocene tectonic activity is likely to have provided the opportunity for a synchronous expansion in the three endemic mammal species in this study, as well as numerous other species. Interestingly, both our Pleistocene geological reconstruction and our proposed origins of expansion in the centre of the island closely resemble maps inferred from a study of tarsier species distribution on Sulawesi [53].

Furthermore, the recent emergence of connections between Sulawesi’s arms coincides with a faunal turnover on the island and the extinction of multiple species. The geological reconstruction, and in particular the recent elimination of the marine barrier at the Tempe depression separating the Southwest and Central regions, fits well with suggested replacement in tarsier species that occurred in the last ~1 My [41]. The extinction of a numerous other species found in the early to late Pleistocene (~2.5–0.8 Myr ago) palaeontological assemblages of the Southwest arm, such as dwarf elephants (*Stegodon sompoensis* and *Elephas celebensis;* [19]), also coincides with the emergence of connections with the Central region. The dispersal of our three species from the central region of Sulawesi may therefore have played a role in these extinctions.

Sulawesi’s development by emergence and coalescence of islands had a significant impact on the population structure and intraspecific morphological differentiation of Sulawesi’s three largest mammals and many other endemic taxa. Thus, while most of Sulawesi’s extant fauna arrived relatively recently, the more ancient geological history of the island (collision of multiple fragments) might have also affected patterns of endemism. Many aspects of Sulawesi’s interconnected natural and geological histories remain unresolved. Integrative approaches that combine biological and geological data sets are therefore essential for reconstructing a comprehensive evolutionary history of Wallace’s most anomalous island.

## Acknowledgments

All data associated with this manuscript are available on Dryad (DOI TBD). We thank Joshua Schraiber and Erik Meijaard for valuable comments. L.A.F.F., J.H, A.L., A. H-B and G.L. were supported by a European Research Council grant (ERC-2013-StG-337574-UNDEAD) and Natural Environmental Research Council grants (NE/K005243/1 and NE/K003259/1). L.A.F.F. was supported by a Junior Research Fellowship (Wolfson College, University of Oxford). P.G. S.G., J.v.d.H, C.A. and D.O. were supported by Flemish government structural funding. A.R was supported by a Marie Curie Initial Training Network (BEAN—Bridging the European and Anatolian Neolithic, GA no. 289966) awarded to M.G.T. M.G.T. is supported by a Wellcome Trust Senior Research Fellowship (GA no. 100719/Z/12/Z). B.J.E was supported by the Natural Science and Engineering Research Council of Canada. This work received additional support from the University of Edinburgh Development Trust, the Roslin Institute, the Balloch Trust and the Stichting Dierentuin Helpen (Consortium of Dutch Zoos). Additional support was also provided by The Rufford Small Grant, Royal Geographical Society, London, the Royal Zoological Society of Scotland and The University of Edinburgh Birrell-Gray Travel Award. We also thank the National Museums of Scotland for logistic support, and the Negaunee Foundation for their continued support of a curatorial preparator. We are also indebted to the Indonesian Ministry of Forestry, Jakarta (PHKA), Sulawesi’s Provincial Forestry Departments (BKSDA); the Indonesian Institute of Science (LIPI); Museum of Zoology, Research Center for Biology, Cibinong (LIPI); and the project’s longstanding Indonesian sponsor, Ir. Harayanto MS, Bogor Agricultural University (IPB) for sample collection/permission.

